# Helix formation during the coupled binding and folding of intrinsically disordered proteins monitored by synchrotron-radiation circular dichroism spectroscopy

**DOI:** 10.1101/640599

**Authors:** Elin Karlsson, Eva Andersson, Nykola C. Jones, Søren Vrønning Hoffmann, Per Jemth, Magnus Kjaergaard

## Abstract

Intrinsically disordered proteins organize interaction networks in the cell in many regulation and signalling processes. These proteins often gain structure upon binding to their target proteins in multi-step reactions involving the formation of both secondary and tertiary structure. To understand the interactions of disordered proteins, we need to understand the mechanisms of these coupled folding and binding reactions. We studied helix formation in the binding of the molten globule-like nuclear coactivator binding domain (NCBD) and the disordered interaction domain from activator of thyroid hormone and retinoid receptors (ACTR). We demonstrate that helix formation in a rapid binding reaction can be followed by stopped flow synchrotron-radiation circular dichroism spectroscopy, and describe the design of such a beamline. Fluorescence-monitored binding experiments of ACTR and NCBD display several kinetic phases including one concentration-independent phase, which is consistent with an intermediate stabilized at high ionic strength. Time resolved circular dichroism experiments show that almost all helicity is formed upon initial association of the proteins, or separated from the encounter complex by only a small energy barrier. Through simulation of mechanistic models, we show that the intermediate observed at high ionic strength likely involves a structural rearrangement with minor overall changes in helicity. Our experiments provide a benchmark for simulations of coupled binding reactions and demonstrate the feasibility of using synchrotron radiation circular dichroism for mechanistic studies of protein-protein interactions.

## Introduction

Intrinsically disordered proteins (IDPs) are abundant in eukaryotic proteomes and govern key cellular processes such as cell cycle regulation, cell growth and development^1^. These proteins contribute to the complexity of the interaction networks within cells by interacting with several ligands, acting as scaffold proteins and presenting accessible sites for post-translational modifications. In total, these mechanisms allow IDPs to form complexes that are responsive to cellular cues^2^. The functions of IDPs often require them to bind specifically to other proteins, but the specificity does not stem from a fixed tertiary structure as in folded proteins. To understand the specificity of IDP interactions, it is necessary to understand how they recognize their partners^3,4^.

Binding reactions of IDPs involve at least two processes: binding and folding. This prompts the question of what occurs first, binding or folding? This question resembles the two paradigmatic models of binding reactions developed to describe substrate recognition in enzymes. In the conformation selection model, the structural change is required for binding, which means that the protein folds independently and then binds to its partner. In the induced fit model, the protein binds in a disordered state and folds while remaining bound^5^. IDPs are unlikely to adopt a tertiary conformation identical to the complex before binding. However, they often sample native-like conformations in the disordered state and transient helices are particularly common. To understand coupled binding and folding mechanistically, we have to describe when in the reaction secondary structure and stabilizing inter-molecular interactions are formed.

IDPs rarely bind exclusively through induced fit or conformational selection. More likely, the binding reactions follow parallel pathways, where the flux through the pathways may vary with the conditions. The structural changes that accompany coupled folding and binding are much larger than for substrate binding. They often involve association of multiple independent segments and formation of secondary structure in both partners. Coupled folding and binding reactions are thus inherently multistate, and the reaction often involves several intermediates with varying degrees of native or non-native^6,7^ secondary and tertiary structure. Kinetic studies can in principle resolve intermediates as their formation and breakdown will result in distinct kinetic phases, but in practice this is not always the case. The ability to resolve different kinetic phases depends both on suitable probes for the reaction steps and the time resolution of the experiment^8^. Therefore, technical limitations often prevent decomposition of the binding pathway into multiple steps. As in protein folding, many reactions thus appear as simple two-state reactions where intermediates are not detected experimentally^9^. To maximize mechanistic information, a reaction mechanism should therefore ideally be described using several observables that report on different types of structure formation.

One of the best studied coupled folding and binding reactions is that between the nuclear coactivator binding domain (NCBD) from CREB binding protein and the interaction domain from activator of thyroid receptors (ACTR). In the complex, NCBD forms a three-helix bundle, around which ACTR wraps three short helices without intra-molecular tertiary contacts (Figure 1)^10^. In the unbound state, ACTR is highly disordered but has transient helicity in the first helix of the complex^11,12^. Free NCBD is usually described as a molten globule^13–15^, but has a propensity to form a partially ordered structure resembling the complex with ACTR^16,17^. The binding reaction thus involves association of the proteins, formation of helices in ACTR, formation of inter-molecular interactions, and rigidification of the helices and tertiary structure of NCBD.

**Figure 1.**
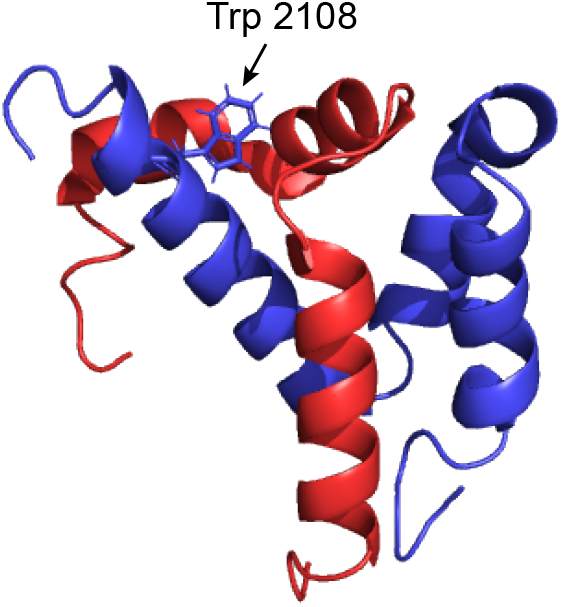
The NMR structure of NCBD (blue) in complex with ACTR (red) with the Trp probe in NCBD highlighted. The figure was created in PyMOL using PDB entry 1KBH^10^

Kinetic experiments are needed to extract information about reaction mechanisms^18^. The reaction between ACTR and NCBD has been studied by stopped-flow fluorimetry and single molecule FRET methods, which showed that the reaction contains multiple intermediates that give rise to observable kinetic phases^8,19–21^. Three kinetic phases can be resolved in the time-scale of stopped-flow measurements: a concentration-dependent phase, which occurs on a shorter timescale than the other phases under our experimental conditions and is hence termed “fast” (λ^fast^). It is mainly determined by the association rate constant, *k_on_*. Furthermore, a concentration-independent phase can be detected under certain conditions over an “intermediate” timescale (λ^intermediate^, τ ≈ 30 ms at 4°C). Finally, a slower concentration-independent phase related to an unknown conformational change is observed on the second timescale (λ^slow^, τ ≈ 1 s at 4°C)^8^. The transition state of the initial association is largely disordered, although the first helix of ACTR is partially formed^19,22^. Single-molecule measurements suggest that additional slower phases occur due to proline cis-trans isomerization^20^. The binding reaction between NCBD and ACTR has been the subject of multiple simulation studies using both coarse-grained and atomistic models^23–26^. These simulations suggest that various intermediates occur on the reaction pathway with varying degrees of native-like secondary structure. It is currently not clear whether the intermediates observed in kinetic studies and in simulations are the same species.

Kinetic studies of folding-and-binding reactions usually rely on fluorescence changes as the detectable signal. Fluorescence has a large signal-to-noise ratio, and many proteins have suitable aromatic residues in binding interfaces that change fluorescence properties upon binding. For proteins without aromatic amino acids in the interfaces, it is often possible to engineer a suitable reporter by conjugation to an extrinsic dye^27–29^ or introduction of a tryptophan residue as e.g. done for NCBD^8^. Fluorescence reports on the local environment around the fluorophore and is thus insensitive to structural changes occurring in the rest of the protein. Furthermore, the fluorescence intensity changes are combined effects of solvation and quenching interactions around the fluorophore. The fluorescence intensity change of partial reactions are thus not necessarily reliable reporters of structural changes. It would therefore be desirable to have other time-resolved observables with a different dependence on structural changes.

Far-UV circular dichroism (CD) spectroscopy reports on secondary structure changes in proteins. Stopped-flow CD can report on secondary structure formation during e.g. a protein folding reaction. Compared to fluorescence detection, CD spectroscopy is much less sensitive and typically requires a 10-fold higher protein concentration. This may be feasible for kinetic studies on folding of monomeric proteins as the folding rate is concentration-independent. Bimolecular binding reactions, however, are usually concentration-dependent. A 10-fold increase of the concentration of one binding partner, thus results in a 10-fold increase in the initial reaction rate. Thus, an increase in the concentration moves the reaction outside the range that can be captured with the stopped-flow technique. This problem can in principle be solved by increasing the sensitivity of stopped-flow CD. For static measurements, synchrotron radiation CD (SRCD) has been used to increase the intensity of the light source, which extends the CD spectrum to lower wavelengths, and permits the use of lower sample concentrations^30^.

Here we report the development of a stopped-flow CD setup coupled to a high intensity UV beam line. We use this setup to follow the formation of helical structure in the coupled folding and binding reaction between ACTR and NCBD. We show that helix formation is coupled to the initial association and that slower kinetic phases likely arise from subtle rearrangements of helical complexes. To our knowledge, this is the first application of time-resolved SRCD to coupled folding and binding reactions, and suggests that the approach can be used to study structural changes accompanying many other rapid binding reactions.

## Materials and methods

### Protein expression and purification

The expression plasmids carried the cDNA sequences of the interaction domain from human ACTR (corresponding to amino acid residues 1023-1093 in UniProt ID Q9Y6Q9-1) and NCBD from human CREBBP (residues 2058-2116 in UniProt ID Q92793-1). The expression constructs were N-terminally tagged with a 6xHis-lipoyl domain. The plasmids were transformed into One Shot BL21(DE3)pLysS Chemically Competent *E. coli* cells (Invitrogen) and selected on LB agar with 100 μg/mL ampicillin and 35 μg/mL chloramphenicol. The cells were grown in LB media with 50 μg/mL ampicillin at 37 °C while shaking until OD600 reached a value of around 0.6. After addition of 1 mM isopropyl ß-D-1-thiogalactopyanoside (IPTG), the proteins were expressed overnight at 18 °C while shaking. The cells were lysed by sonication and the lysate was separated on a Ni sepharose 6 Fast Flow resin (GE Healthcare) equilibrated with 30 mM Tris-HCl pH 8, 500 mM NaCl as the binding buffer and 30 mM Tris-HCl pH 8, 500 mM NaCl, 250 mM imidazole as the elution buffer. After cleavage of the 6xHis-lipoyl domain with thrombin (GE Healthcare), the protein was subjected to separation on the Ni sepharose 6 Fast Flow resin (GE Healthcare) using the same buffers as above to separate the protein from the cleaved tag. The proteins were further separated on a C8 Reversed phase column (Vydac), using a 0-70 % acetonitrile gradient and fractions containing pure proteins were lyophilized. The fractions were analysed with MALDI-TOF mass spectrometry to ensure the correct identity. The concentrations of the proteins were measured by A280 for proteins containing Trp or Tyr residues. The extinction coefficients were calculated using the ExPASy ProtParam online software^31^. ACTR, which is lacking aromatic residues, was measured by A205 and the concentration was estimated using an extinction coefficient 250,000 M^-1^ cm^-1^.

### Static SRCD spectroscopy

Lyophilized protein was dissolved in 20 mM sodium phosphate buffer pH 7.4 with 150 mM NaF to reduce buffer absorbance at low wavelengths. The protein concentrations were 69 μM ACTR and 80 μM NCBD, which gives a maximum in absorbance of approximately 1 in a 0.1 mm quartz cuvette. The exact pathlength of the cuvette was determined by an interference technique to be 0.1027 mm^32^. The spectra were recorded on the AU-CD beam line, ASTRID2, at 25°C in three scans between 280 and 170 nm. A reference baseline spectrum of the buffer alone was subtracted, and all three spectra were averaged and mildly smoothed with a 7 point Savitzky-Golay filter^33^.

### The rSRCD facility used for stopped-flow SRCD measurements

The rapid Synchrotron Radiation Circular Dichroism (rSRCD) facility is located on the AU-rSRCD branch line of the AU-AMO beam line at the synchrotron radiation facility ASTRID2, at Aarhus University in Denmark. Figure 2 shows a schematic overview of the beam line. The source of this beam line has a 2.4 m long undulator with a magnetic period of 53 mm, which, in combination with the relatively low electron energy of the ASTRID2 accelerator of 0.58 GeV, allows the generation of intense UV light up to a wavelength of 290 nm. The small divergence and narrow width of this UV light source makes it well suited for a stopped flow instrument, where the internal light beam width must be kept below 1 mm. This can be achieved without any further optics than those in the AU-AMO monochromator which selects the UV wavelength of interest. Note that this wavelength can be freely chosen below 300 nm, and is thus not restricted as more conventional sources are, where intense single lines of a lamp limits the wavelength to certain discrete values. The UV light from the AU-AMO monochromator is directed into the rSRCD branch line via a flat MgF_2_ protected aluminium coated mirror. This mirror has a sideways, as well as a reflection angle adjustment, which makes easy alignment of the light into the stopped flow instrument.

**Figure 2.**
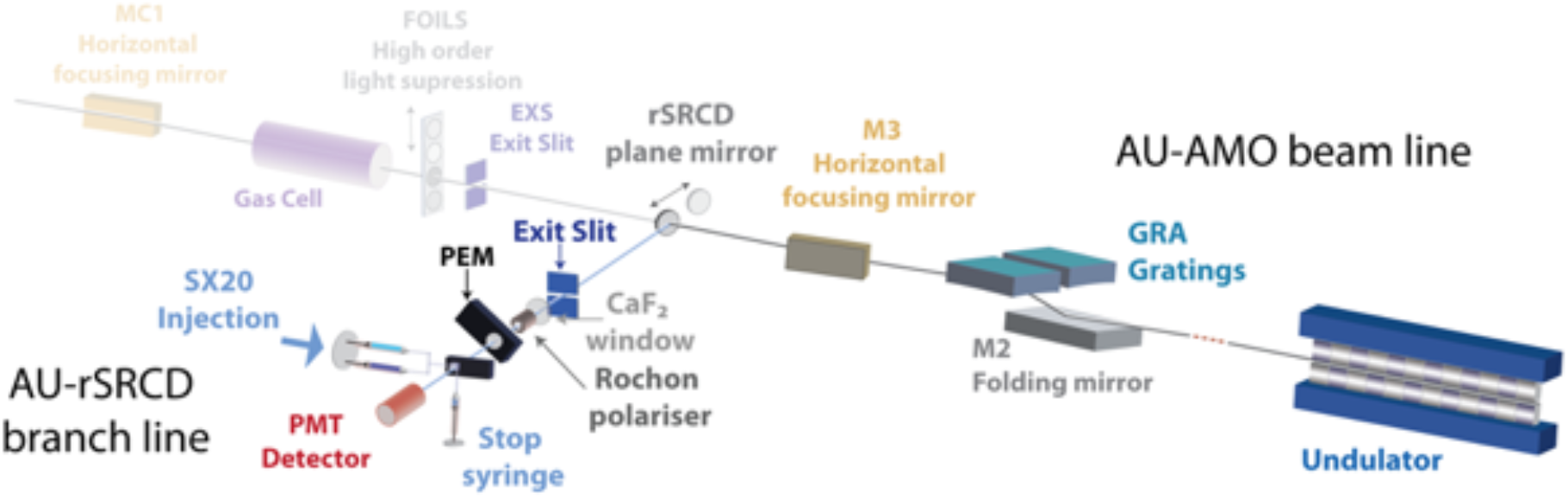
Overview of the AU-AMO beam line and rSRCD branch line at the synchrotron radiation facility ASTRID2, Aarhus University, Denmark. The beam is extracted from the existing beam line (AU-AMO) using a moveable plane mirror, and connected to a commercial SX20 stopped-flow module.

After the exit slit in the rSRCD branch line, which selects the wavelength from the grating in the monochromator, the light passes through a high quality excimer laser graded CaF2 window (Vacom, Germany). The window is cut perpendicular to the <111> crystal axis of CaF2 to minimize birefringence, which would otherwise change the polarization of the light. This window separates the ultra-high vacuum of the AU-AMO beam line from the nitrogen-purged environment of the stopped flow system.

The polarization of the light from the monochromator is nearly fully horizontally polarized, but to ensure 100% linear polarization, the light passes through a MgF_2_ Rochon polarizer (B.Halle GmbH, Berlin, Germany). The following optical element, a Photo Elastic Modulator (PEM – Hinds, Oregon, USA), converts the horizontally polarized light into alternating left and right handed circularly polarized light with a frequency close to 50 kHz. Thus, the CD signal of a chiral sample will oscillate in phase with this frequency, and can be detected using a lock-in amplifier.

The stopped flow instrument chosen for the rSRCD facility is a model SX20 from Applied Photophysics Limited (APL – Surry, UK), which can perform multiple fully automated mixing shots. The photomultiplier tube (PMT) and data acquisition electronics from APL records CD signal, with time resolutions down to 0.1 ms. The APL electronics controls both the PMT high voltage and the PEM, using the DataPro SX20 software. The wavelength of the beam line is set via the ASTRID2 control system software (ConSys), which can set both the monochromator and the undulator to the desired wavelength, as well as directly access the APL software for setting the PEM to the correct setting for CD measurements.

The stopped flow cell in the SX20 stopped flow system is a clear quartz cell; mixing of dark and clear quartz, often used in fluorescence and UV-VIS absorption stopped flow instruments, is avoided to minimize stress-induced birefringence in the quartz. This would otherwise interfere with the circular polarization state of the light, and thus might lead to incorrect CD measurement. Careful masking of the clear quartz avoids light passing through the quartz, and thus not through the sample, to reach the PMT detector, which otherwise can led to a falsely low apparent CD signal of the sample. The quartz cell can be oriented in two directions, with either 2 mm or 10 mm light pathlength. The former is used for most experiments. The two sample syringes and the quartz cell can be submerged in a water bath to set the sample temperature using a water cooler/heater. Typically the temperature can be set between 5°C and 50°C, with temperature oscillations of about 0.1°C. The two mixing syringes of the SX20 system can readily be exchanged for either two 1 mL syringes or one 2.5 mL and one 0.25 mL syringes, thus providing either 1:1 or 1:10 mixing.

### Calibration of the T0 of mixing of the rSRCD setup

To calibrate the deadtime of the instrument, we studied the reaction between N-acetyltryptophane amide (NATA) with N-bromosuccimide (NBS). This reaction is typically used to calibrate fluorescence detecting stopped flow instruments because it follows simple bimolecular kinetics and can be monitored optically due to the quenching of NATA fluorescence upon NBS binding. In the present rSRCD setup, NATA/NBS reactions were followed by observation of the absorbance at 280 nm. Reactions were recorded for final concentrations of 20 μM NATA and NBS ranging from 100-1250 μM at 6 °C. The kinetic traces were fitted to a single exponential function and the intersection of the fitted lines were taken as the real time zero in the experiment, which is 3 ms before the start of the data collection.

### ACTR:NCBD binding monitored by rSRCD

The stopped flow experiments were conducted in 20 mM sodium phosphate buffer pH 7.4 with 150 mM NaF (low ionic strength conditions, *I* ≈ 0.20 M) or with 0.90 M NaF (high ionic strength conditions, *I* ≈ 0.95 M). All experiments were performed at 5°C. In all experiments, the concentration of NCBD_Y2108W_ was held constant at 2.0 μM and the ACTR concentration was varied between 1.7-6.9 μM. The analyzed kinetic traces were typically an average of >20 individual traces. The binding reaction was monitored with the CD signal at 195 nm.

### Stopped flow fluorescence spectroscopy

The fluorescence-monitored stopped flow experiments were carried out on an upgraded SX-17MV stopped flow-spectrometer (Applied Photophysics). The excitation wavelength was set to 280 nm and the emitted light was passed through a 320 nm cut-off filter prior to detection. The buffer conditions and temperature were the same as for the stopped flow CD measurements. The displacement experiments, in which *k_off_^app^* was determined, were conducted using a premixed solution of 1 μM NCBD_Y2108W_ and 0.5 μM WT ACTR (~ 0.5 μM complex) and using an excess of WT NCBD (up to 30-fold) to displace NCBD_Y2108W_ from the complex. Each analyzed kinetic trace was typically an average of 6-8 individual traces. The real time zero was determined by mixing NATA with excess of NBS and monitoring the fluorescence emission. The kinetic traces were fitted to a single exponential function and the intersection of the fitted lines was used as an estimate of the real time zero in the experiment, which was at −1.25 ms in relation to the first recorded data points.

### Data analysis and fitting

The stopped flow kinetic traces were fitted globally by numerical integration using KinTek Explorer (KinTek Corporation) in order to extract values of the microscopic rate constants for the different models^34,35^. Confidence contour plots were computed to assess how constrained the parameters were for each fit. The confidence contour plots show the variation in the χ^2^ ratio (χ^2^_min_/χ^2^) as each parameter is varied individually while all other parameters are kept constant. Well-constrained parameters yield confidence contour plots with well-defined χ^2^ ratio maxima. The lower and upper confidence interval bounds were estimated from a (χ^2^_min_/χ^2^) ratio of 0.9, according to the recommendations in the KinTek user manual (https://www.kintekexplorer.com/docs/Kintek_Explorer_Instructions.pdf). The real time zero used for analysis was −1.25 ms for the fluorescence-monitored stopped flow experiments and – 3 ms for the CD-monitored stopped flow experiments. The total mixing time (or dead time) in the respective stopped flow experiment was 3.25 ms and 6 ms, respectively. This is the time between the real time zero and the first data points, which fall on the fitted single exponential curve. The ACTR/NCBD binding traces were fitted using scaling factors (multiplier or offset for fluorescence and CD data, respectively) to account for small fluctuations in lamp intensity and errors in concentration in the fluorescence-monitored measurements and offset intercept values in the CD-monitored data. Individual traces were fitted to either a single-(Equation 1) or a double-exponential function (Equation 2) in GraphPad Prism 6.0 (GraphPad Software).

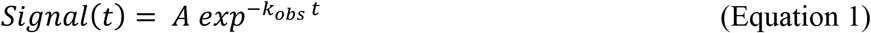

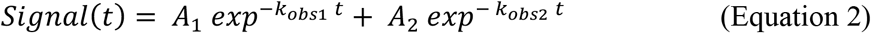

Where A is the amplitude of the kinetic phase, *k_obs_* is the observed rate constant and *t* is time.

## Results

### Test of the SRCD operation using a standard sample

The circular dichroism performance of both the static and stopped flow systems was checked by scanning a camphor sulfonic acid (CSA) sample between 190 nm and 300 nm. The 290.5 nm (positive) CD peak of CSA was used to calibrate the CD signal magnitude, while the 192 nm (negative) peak to 290.5 nm peak ratio was used to judge birefringence of the quartz cell. The first cell delivered with the SX20 instrument, showed a ratio of close to 1, which is far from the 2.1 ratio often quoted for CSA^36^. A screening of quartz cells performed by the manufacturer identified a cell with a CSA ratio close to 2, which is now in use at the beam line. The stopped flow system on the rSRCD facility is therefore calibrated and documented down to 190 nm.

### SRCD allows helix formation to be followed using lower wavelength bands

The coil-to-helix transition in ACTR upon binding to NCBD makes this reaction suited for CD. Previously, it was found that binding resulted in a large change in CD signal at 222nm^10^. We repeated these measurements using the static SRCD beamline, which allowed us to extend the spectrum to shorter wavelengths (Figure 3). The spectral change in the reaction can be estimated by comparing the spectrum of the complex and the summed spectra of the individual proteins. This showed that while the reaction can be followed at 222 nm as is customary in benchtop instruments, a ~3-fold larger change occurs at lower wavelengths. We thus followed the binding reaction at the wavelength with the maximal signal change at 195 nm (Figure 3).

**Figure 3.**
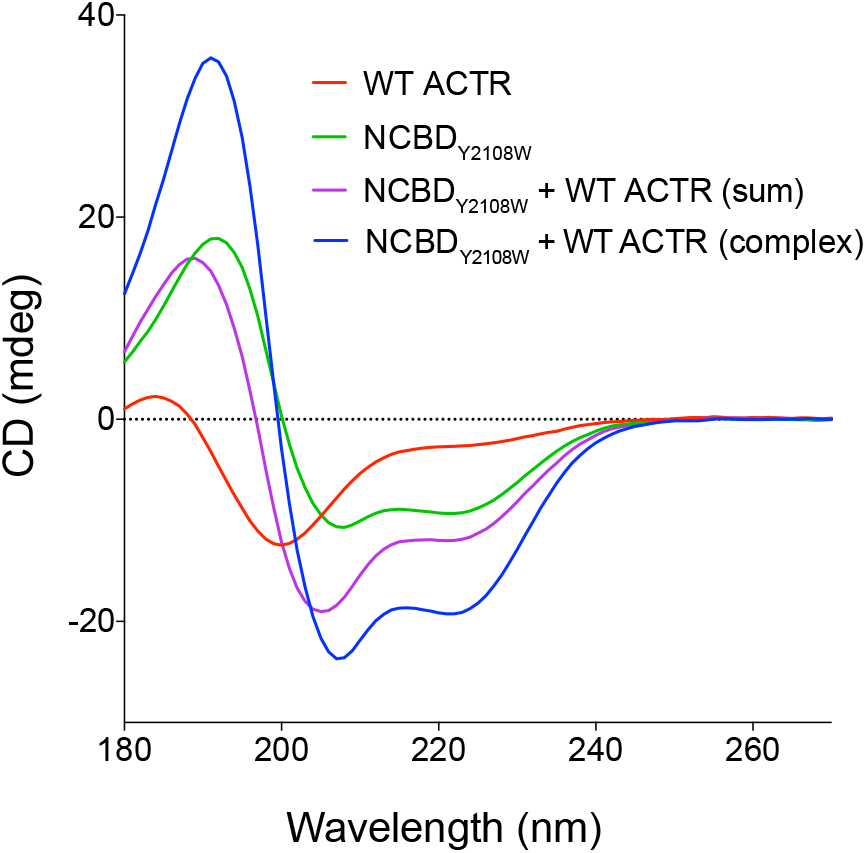
Static CD spectra of free wildtype (WT) ACTR (red) (69 μM), free NCBD_Y2108W_ (green) (80 uM), the theoretical sum of free WT ACTR and free NCBD_Y2108W_ (purple) and WT ACTR (blue) in complex with NCBD_Y2108W_ (69 μM and 80 μM, respectively). The dotted line shows the 0-level of the CD signal.

### Following the association reaction between ACTR and NCBD by rSRCD

To assign helix formation to the different kinetic phases of the binding pathway, we wanted to follow the binding reaction by rSRCD. We mainly performed kinetic experiments using a pseudo-wildtype NCBD_Y2108W_, which enabled fluorescence detection of the reaction in a separate experiment. Previous experiments showed that this mutation does not affect the binding kinetics or secondary structure content of NCBD^8^. The experimental conditions involve several trade-offs related to the protein concentrations that will likely be general to rSRCD experiments of coupled folding and binding. Increased protein concentration improves the signal, but also increases the absorption and thus the noise. Large excess of one binding partner simplifies the reaction to a pseudo-first order reaction, but also increases the reaction rate and shifts part of the signal amplitude into the deadtime. We mixed NCBD with a small excess of ACTR and observed the CD signal at 195 nm. The binding reaction gave rise to a detectable exponential decay. However, much of the reaction amplitude is lost due to a large artefactual signal such that the first 3 ms of the time trace had to be discarded (in addition to the 3 ms of the reaction occurring before the instrument starts recording data). The absolute CD signal displayed significant baseline shift from one acquisition to the next. This persisted in pure water, and we believe both of these artefacts arise due to small distortions of the cuvette when it is pressurized by the stopped-flow setup. Despite these limitations, we managed to obtain kinetic time traces for this binding reaction across a concentration range for the ligand in excess, such that *k_on_* could be determined.

### Binding follows an apparent two-state mechanism at physiological ionic strength

The reaction between ACTR and NCBD has been previously studied by stopped flow fluorimetry^8,37,38^. The current rSRCD measurements were, however, conducted under different buffer conditions to reduce the absorption of buffer components at low wavelengths. To test whether the change in buffer conditions altered the reaction, we re-recorded kinetics using stopped-flow fluorimetry under identical conditions as used for the rSRCD experiments. The binding reaction was first measured at physiological ionic strength conditions (≈ 0.20 M). Under these conditions, the fluorescence-monitored stopped flow traces were monophasic (Figure 4a) with a rate constant that increased linearly with increasing ACTR concentration and thus report on the bimolecular association. Previously, a slow (τ ≈ 1 s at 4°C) concentration-independent phase, λ^slow^, was observed at physiological ionic strength^8^. We did not probe this slow kinetic phase, because rSRCD measurements resulted in radiation-damage of the protein on the timescale where it occurs (1-10 s).

**Figure 4.**
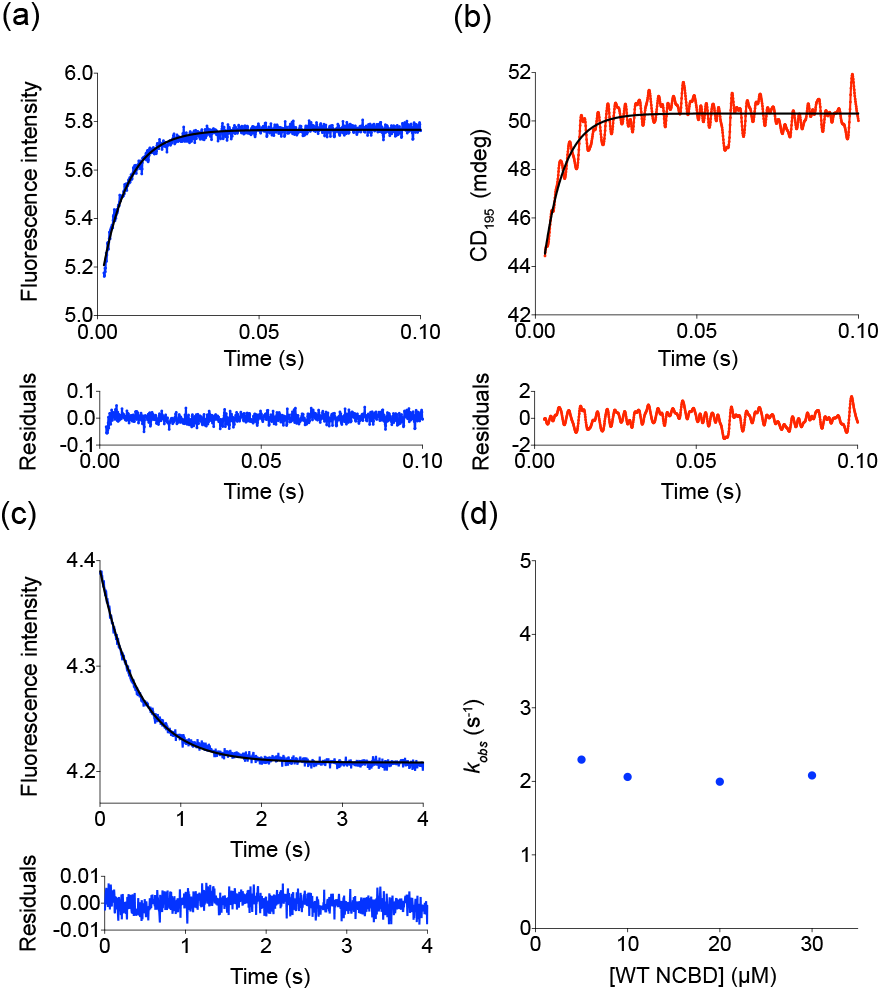
Stopped flow kinetic traces of pseudo-wildtype NCBD_Y2108W_ and ACTR monitoring fluorescence intensity (blue) and CD_195_ (red) at low ionic strength conditions (20 mM sodium phosphate buffer pH 7.4, 150 mM NaF) at 5°C. Kinetic traces for 2.0 μM NCBD mixed with 5.2 μM ACTR are shown in a-b. (a) The fast kinetic phase observed in the fluorescence-monitored data was fitted to a single exponential and yielded a k_obs_ value of 124 ± 1 s^-1^. (b) The CD-monitored binding trace was fitted to a single exponential function with a k_obs_ value of 157 ± 5 s^-1^. (c) A typical displacement experiment monitored by fluorescence where 30 μM wildtype NCBD was used to displace NCBD_Y2108W_ from its complex with ACTR (≈ 0.5 μM complex). The kinetic traces were fitted to a single exponential function and the residuals for the fit are shown below the plot. (d) The kinetic traces from displacement were fitted and k_obs_ values plotted against the concentration of wildtype NCBD. The average of the k_obs_ values at the three highest concentrations were taken as an estimate of k_off_ and the standard deviation was taken as the error, resulting in an estimated k_off_ = 2.05 ± 0.05 s^-1^.

To monitor helix formation in the binding reaction, we first performed rSRCD experiments at physiological ionic strength. The reaction transient monitored via CD_195_ could be described by a single exponential (Figure 4b) with a rate constant linearly increasing with concentration. We obtained the association rate constant, *k_on_* by varying the concentrations of ACTR and fitting the time traces globally to a two-state binding reaction using a numerical integration method. We note that the *k_off_* value obtained from the CD-monitored data was substantially higher compared to the fluorescence-monitored data. However, this is likely a result of the large experimental noise in the stopped flow CD data, which causes an uncertainty in the globally fitted *k_off_* value. Accordingly, confidence contour analysis revealed that *k_off_* was not well-constrained in the global fitting. Thus, *k_off_* is best determined in a separate displacement experiment (Figure 4c). Using fluorescence-monitored stopped flow, the displacement experiment yielded an estimated *k_off_* of 2.05 ± 0.05 s^-1^ (Figure 4d). The fitted value for *k_on_* was similar for the CD- and fluorescence-monitored binding reactions (Table 1) suggesting that fluorescence and CD signals report on the same apparent two-state transition. This shows that helix formation and binding are likely coupled or only separated by a low energy-barrier, which prevents these two steps from being resolved. In addition, we found that the NCBD_Y2108W_ pseudo-wildtype exhibited almost identical binding kinetics to wildtype NCBD when the reaction was monitored with the CD_195_ signal, which confirms that the Trp mutation does not influence the mechanism of interaction^8^ (Table 1).

**Table 1.** Fitted rate constants for the binding reactions of NCBD and ACTR. The conditions were 20 mM sodium phosphate buffer pH 7.4, 150 mM NaF and the stopped flow experiments were performed at 5°C. The association rate constants (*k_on_*) were obtained from fitting the stopped flow time traces globally by numerical integration to a two-state binding mechanism. The confidence interval boundaries were calculated using a χ^2^ ratio threshold of 0.9 in all cases. *The *k_off_* value was determined in a separate fluorescence-monitored displacement experiment. Wildtype NCBD was used in large excess to displace NCBD_Y2108W_ from the complex with ACTR. The value of *k_off_* is the average from the three highest WT NCBD concentrations and the error is the standard deviation of these measurements.

| NCBD variant | ACTR variant | Parameter | Best fit value | Lower bound | Upper bound | Method |
| --- | --- | --- | --- | --- | --- | --- |
| NCBD<sub>Y2108W</sub> | WT ACTR | $k_{on}$ | $27 \mu\text{M}^{-1} \text{s}^{-1}$ | 17 | 42 | SR-CD stopped flow |
| | | $k_{on}$ | $30 \mu\text{M}^{-1} \text{s}^{-1}$ | 28 | 33 | Fluorescence-monitored stopped flow |
| | | $k_{off}^*$ | $2.05 \pm 0.05 \text{s}^{-1}$ | - | - | Fluorescence-monitored stopped flow |
| WT NCBD | WT ACTR | $k_{on}$ | $28 \mu\text{M}^{-1} \text{s}^{-1}$ | 18 | 40 | SR-CD stopped flow |

**Table 2.** Fitted rate constants for binding reactions of NCBD_Y2108W_ and WT ACTR. The stopped flow experiments were performed in 20 mM sodium phosphate buffer pH 7.4, 0.90 M NaF at 5°C. The rate constants were obtained by fitting fluorescence-monitored stopped flow data to either an induced fit mechanism or a conformational selection mechanism using a numerical integration software. The best fit value of each parameter is shown along with the lower and upper confidence interval bounds. The confidence intervals bounds were computed using a χ^2^ ratio threshold of 0.9 for both models.

| Model | Parameter | Best fit value | Lower bound | Upper bound |
| --- | --- | --- | --- | --- |
| Induced fit | $k_{+1}$ | $33 \mu\text{M}^{-1} \text{s}^{-1}$ | 26 | 41 |
| | $k_{-1}$ | $9.3 \text{s}^{-1}$ | 3.0 | 20 |
| | $k_{+2}$ | $15 \text{s}^{-1}$ | 3.2 | 22 |
| | $k_{-2}$ | $9.3 \text{s}^{-1}$ | 6.5 | 17 |
|  | Scaling factor 1 | (1.0) | - | - |
|  | Scaling factor 2 | 0.98 | 0.98 | 0.98 |
|  | Scaling factor 3 | 0.98 | 0.98 | 0.99 |
|  | Scaling factor 4 | 0.98 | 0.98 | 0.99 |
|  | Scaling factor 5 | 0.97 | 0.97 | 0.98 |
|  | Scaling factor 6 | 0.97 | 0.97 | 0.98 |
|  | Scaling factor 7 | 0.97 | 0.96 | 0.98 |
| Conformational selection | $k_{+1}$ | $25 \text{s}^{-1}$ | 20 | 31 |
| | $k_{-1}$ | $6.9 \text{s}^{-1}$ | 1.2 | 13 |
| | $k_{+2}$ | $29 \mu\text{M}^{-1} \text{s}^{-1}$ | 27 | 36 |
| | $k_{-2}$ | $1.7 \text{s}^{-1}$ | 1.1 | 2.7 |
|  | Scaling factor 1 | (1.0) | - | - |
|  | Scaling factor 2 | 0.98 | 0.98 | 0.98 |
|  | Scaling factor 3 | 0.99 | 0.99 | 0.99 |
|  | Scaling factor 4 | 0.99 | 0.99 | 0.99 |
|  | Scaling factor 5 | 0.98 | 0.98 | 0.99 |
|  | Scaling factor 6 | 0.98 | 0.98 | 0.99 |
|  | Scaling factor 7 | 0.98 | 0.98 | 0.99 |

### Kinetic experiments at high ionic strength probe an additional reaction step

Previous studies revealed the presence of an additional concentration-independent kinetic phase, λ^intermediate^ (τ ≈ 30 ms at 4°C), at high ionic strength (≈ 0.95 M)^8^. This suggests the presence of a salt-dependent, kinetic intermediate in the binding reaction of NCBD and ACTR. Before repeating these experiments by rSRCD, we performed fluorescence-monitored kinetic measurements of NCBD_Y2108W_ and wildtype ACTR in 0.90 M NaF. The resulting kinetic traces were clearly biphasic, as shown before in presence of 0.90 M NaCl^8^ (Figure 5 a-b). Thus, fluorescence-monitored binding at 0.90 M NaF shows that the interaction between ACTR and NCBD is at least a two-step process described by the kinetic phases λ^fast^ and λ^mtermediate^. As previously, the radiation-damage at longer time scales prevented studying λ^slow^ with rSRCD.

**Figure 5.**
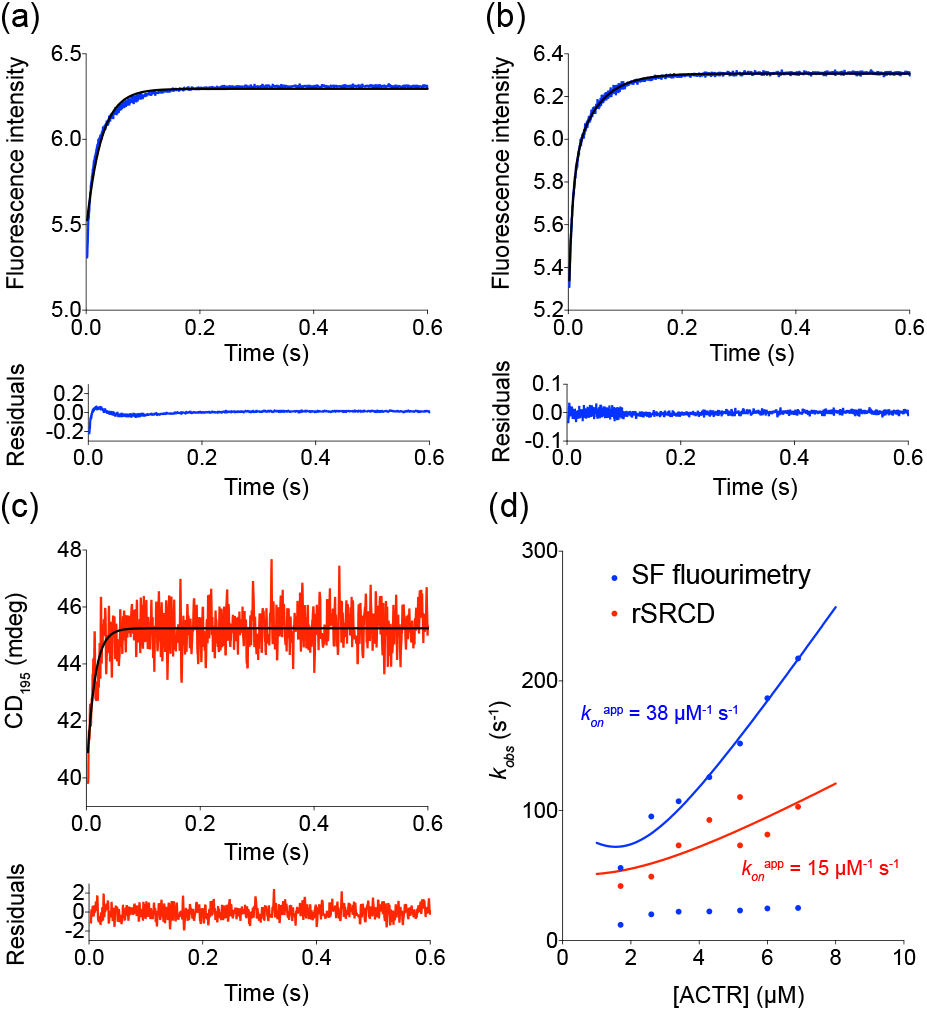
Stopped flow binding traces of pseudo-wildtype NCBD_Y2108W_ and ACTR at high ionic strength conditions (20 mM sodium phosphate buffer pH 7.4, 0.90 M NaF) at 5°C, monitored by fluorescence intensity (blue) or CD_195_ (red). The concentrations used in the experiments were 2.0 μM NCBD and 1.7-6.9 μM ACTR. Examples of traces are shown for 2.0 μM NCBD and 5.2 μM ACTR. (a) The fluorescence-monitored kinetic trace was fitted to a single exponential function and the residual plot shows a clear deviation from a random distribution. (b) The fluorescence-monitored kinetic trace fitted to a double exponential function. The residual plot shows a close to random distribution. The fitted k_obs_ values were 152 ± 2 s^-1^ for λ^fast^ and 23.1 ± 0.2 s^-1^ for χ^mtermediate^. (c) The CD-monitored trace was apparently monophasic and was fitted to a single exponential function yielding an apparent k_obs_ value of 73 ± 3 s^-1^. (d) Plotting the fitted rate constants from the fluorescence-monitored kinetic experiments (blue dots) against concentration of ACTR revealed one concentration-dependent kinetic phase which reports on the binding event and one concentration-independent phase with a k_obs_ of around 20 s^-1^. The fitted rate constant from the CD-monitored experiments (red dots) showed an approximately linear dependence on ACTR concentration, but with a lower apparent k_on_^app^ value. The solid lines represent the fits of the fluorescence and CD-monitored data to an equation for bimolecular association^44^, which was used to extract the k_on_^app^ values reported in this figure. The observed rate constants from the CD-monitored experiments are likely weighted averages of λ^fast^ and λ^intermediate^.

In principle, one could differentiate between induced fit and conformational selection if the fluorescence yields of the molecular species in the reaction were known. As this information is not known, we cannot infer a mechanism from the amplitude information in a fluorescence-monitored experiment. On the other hand, amplitudes obtained in a CD-monitored experiment are directly related to the amount of helicity in the protein. Thus, to probe helix formation in the different steps of the binding reaction, stopped flow measurements at high ionic strength were conducted monitoring the CD_195_. The stopped flow CD traces were apparently monophasic under high ionic strength conditions and no intermediate kinetic phase was observed (Figure 5c). However, plotting the fitted *k_obs_* values against concentration of ACTR showed that the apparent *k_on_* (*k_on_*^app^) from the SR-CD stopped flow experiments was distinctively lower than *k_on_*^app^ from the fluorescence-monitored experiments (Figure 5d). This suggests that the fast and intermediate kinetic phases cannot be resolved at the present noise level. The apparent *k_obs_* value from the fit of a single exponential function to CD_195_ thus constitutes an average of the *k_obs_* values from λ^fast^ and λ^intermediate^ weighted by their fractional amplitude. This likely results in the apparently lower *k_on_*^app^ observed for the CD-monitored stopped flow data. In fact, the same phenomenon was observed in fluorescence-monitored binding experiments when NCBD was in excess over ACTR^39^.

Induced fit and conformational selection have a different order of concentration dependent and independent steps, and can thus occasionally be distinguished from global fits of experimental traces recorded over a concentration series (Figure 6b). The kinetic traces from experiments at high ionic strength were fitted globally by numerical integration to either an induced fit or a conformational selection model to extract the rate constants associated with the steps in the respective three-state model (Figure 6a-b). Based solely on the fluorescence-monitored data, one of these models could not be preferred over the other. The confidence contour analysis suggested that the parameters were more well-constrained for the induced fit model and *k-1* was not well-constrained in the fit to the conformational selection model (Figure 6c). Neither of these two-step models describe the data perfectly, but the deviation from a perfect fit was deemed too small to invoke more steps in the mechanism.

**Figure 6.**
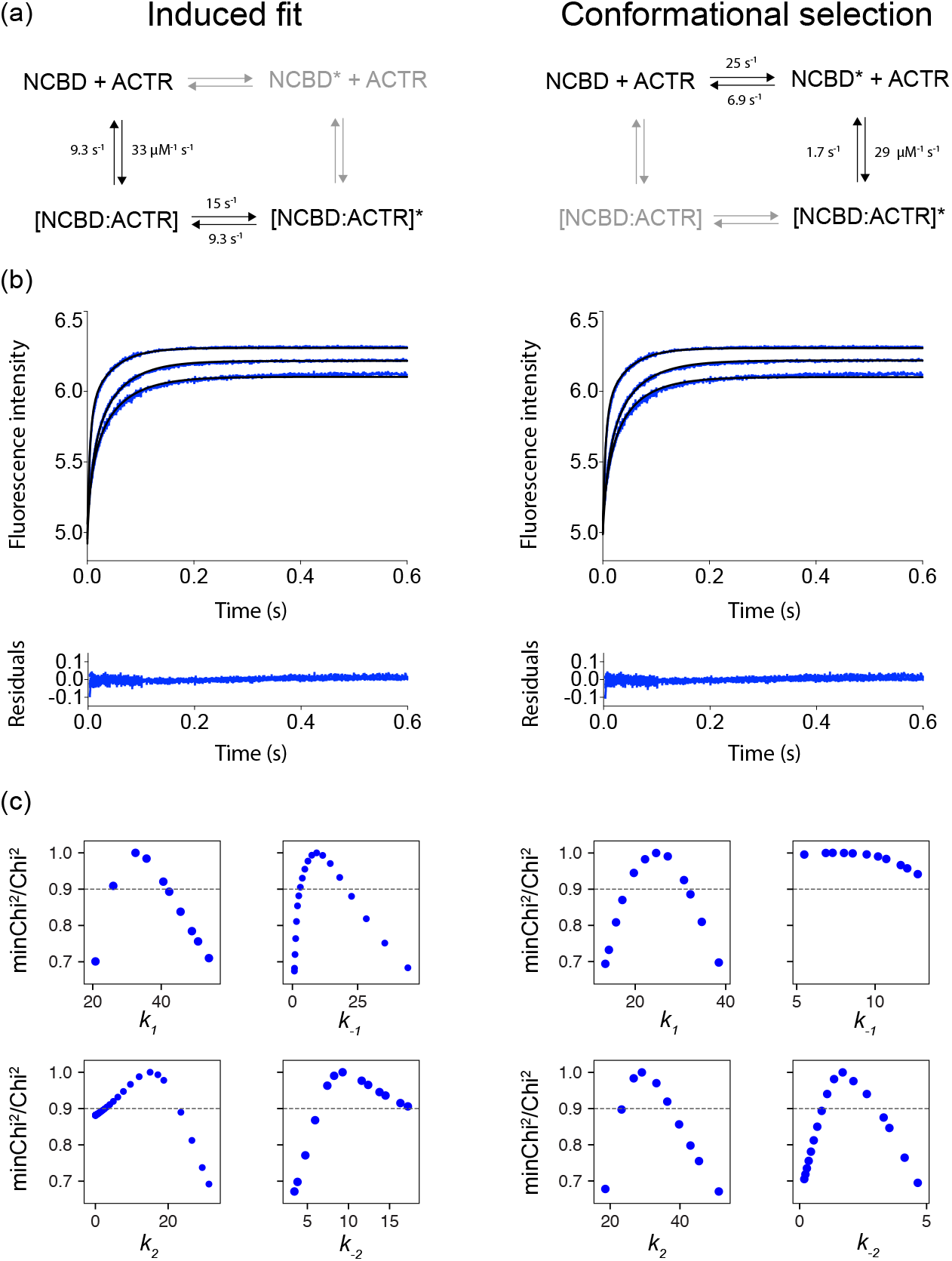
Stopped flow fluorescence-monitored data recorded at high ionic strength conditions (20 mM sodium phosphate buffer pH 7.4, 0.90 M NaF) at 5*C. The concentration of pseudo-wildtype NCBD_Y2108W_ was held constant at 2.0 μM and ACTR was varied between 1.7-6.9 μM. (a-b) The data set was fitted globally by numerical integration to either an induced fit model (left panel) or a conformational selection model (right panel). We use an asterisk to denote a molecular species which has undergone a structural rearrangement. Three out of seven traces are shown in the figure for clarity. The residual plots for each fit is shown below the curve. Based on the residual plot, both models fit the data equally well and best fit parameters are shown in the respective scheme. (c) Confidence contour plots for each fitted model show that the value for the parameter k-1 was poorly constrained in the fit to the conformational selection model, as judged by the absence of a well-defined maximum in the χ^2^_min_/χ^2^ ratio. The dotted grey line shows the χ^2^ ratio threshold of 0.9, which was used to calculate the lower and upper 95 % confidence bounds.

### Kinetic simulations suggest that the intermediate is helical

The time-resolved SRCD experiments did not allow us to extract amplitude information for λ^fast^ and λ^intermediate^ by direct fitting. Therefore, we simulated kinetic profiles for possible scenarios for helix formation in induced fit and conformational selection models, and compared the simulated and experimental *k_obs_* values. In all simulations, we used the rate constants obtained in the fit of the fluorescence data above. We defined three different possible models for helix formation in ACTR upon binding to NCBD (Figure 7a): 1) Induced fit where ACTR has no helical content in the intermediate, 2) Conformational selection where NCBD undergoes a conformational rearrangement with no overall changes in helicity prior to binding to ACTR and 3) Induced fit where ACTR has all helical content in the intermediate. The fourth possible model, conformational selection in ACTR was ruled out since this model resulted in poor fits to the data. The asterisk annotation of some of the molecular species in the reaction schemes indicates structural rearrangement (Figure 7a).

**Figure 7.**
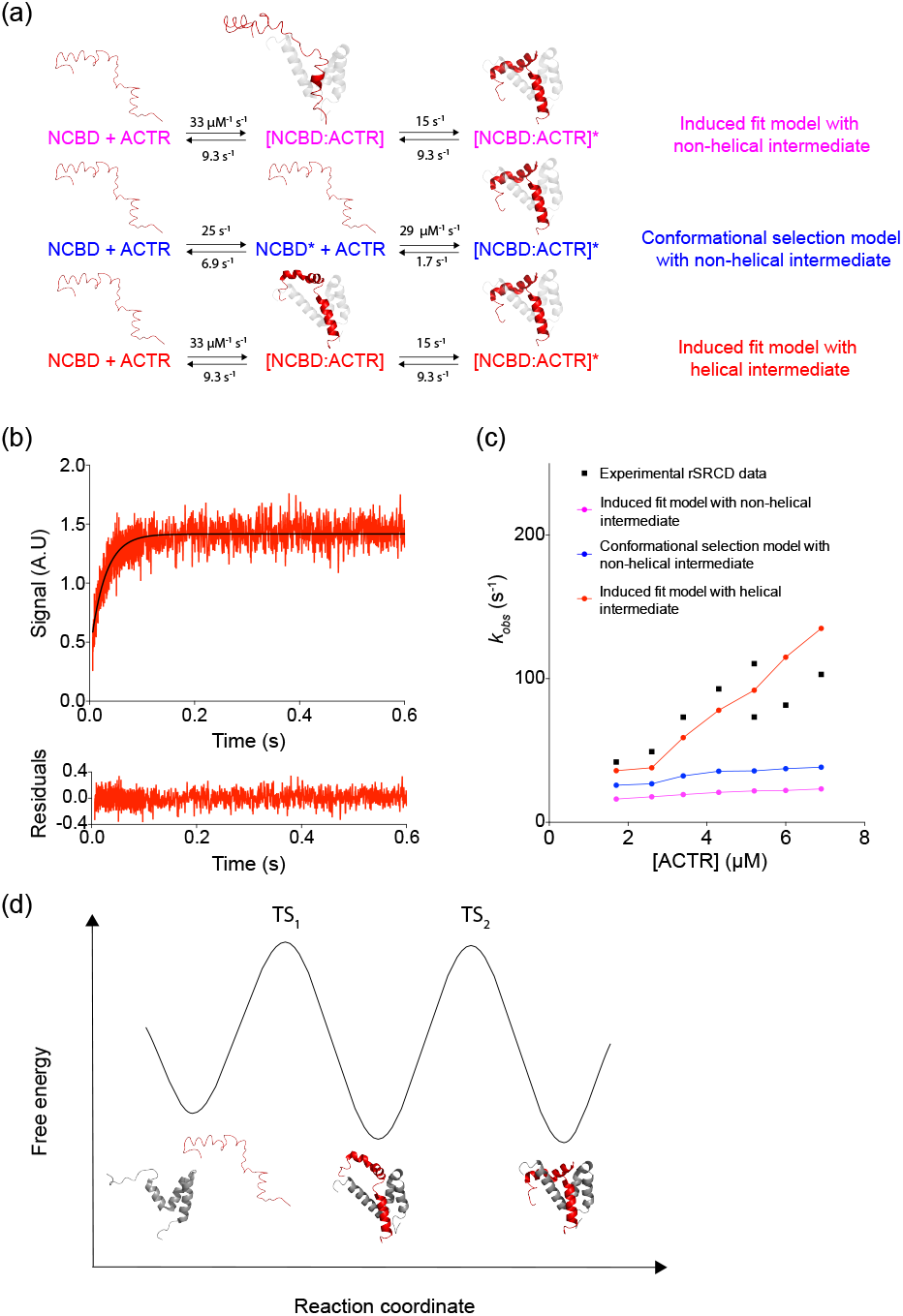
(a) The simplest scenarios for helix formation were defined for different two-step mechanisms. The first model is an induced fit model where all helix formation occurs in the conformational rearrangement step following initial binding. The second model represents a conformational selection mechanism where a conformational rearrangement of NCBD with no overall changes in helicity is followed by a binding step where all helical content of ACTR is formed. The third model is an induced fit model where all helical content is formed in the initial binding step. The output amplitude factors, which links the signal output to the concentration of species, were defined as either 0 (no helix) or 1 (helices fully formed) for these simplified models. The structures of ACTR above the species are representations of the helical state of ACTR (disordered with transient helices formed vs. stable helices formed) in each step of the different models for binding. (b) Binding reaction profiles were simulated for all models, using the rate constants that were obtained from fitting the fluorescence-monitored stopped flow data at high ionic strength conditions. Gaussian noise was added to the simulated reaction profiles with a standard deviation of 0.1. The traces were fitted to a single exponential function and the residual plot is shown below the graph. The figure shows a simulated binding trace for model 3 as an example. The concentrations used to simulate the trace were 2 μM NCBD and 1.7 μM ACTR. (c) The dependence of apparent k_obs_ on [ACTR] for the defined models. The apparent k_obs_ values were obtained from the fits to the simulated reaction profiles for each model. The apparent k_obs_ values obtained in the SR-CD stopped flow experiments are shown in black squares. (d) An energy diagram with a schematic model for the best-fit scenario of helix formation in ACTR. The energy diagram was computed using 10 μM ACTR. ACTR binds to NCBD with an induced fit mechanism where virtually all helical content is formed in the binding step. The subsequent conformational rearrangement step is not associated with any changes in helicity. The first transition state (TS_1_) was probed previously with Φ-analysis using a set of helix-modulating mutants, which showed that helix 1 in ACTR is around 50 % formed in TS_1_^41^. The models for free NCBD and NCBD in complex with ACTR were created using the PDB entries 1KBH and 2KKJ, respectively^10,16^. The model for the intermediate represents a plausible scenario based on the results in this study.

The relative amplitudes for the CD signal in each model were defined based on the degree of helicity in the intermediate and product states, and was either 0 for non-helical or 1 for fully formed helices. To imitate the signal-to-noise level in the rSRCD experiments, (Figure 7b) Gaussian noise with a standard deviation of 0.1 was added to the traces. The resulting traces were apparently monophasic, and we could fit apparent *k_obs_* values that represent a weighted average of the *k_obs_* values of λ^fast^ and λ^intermediate^ (Figure 7c). The simulated data showed that the induced fit model with a non-helical intermediate was expected to be dominated by λ^intermediate^ and *k_obs_* was approximately constant over the entire concentration range. Furthermore, the reaction profiles of this model contained lag phases which were not observed in the experimental data, although they could be hidden in the experimental noise. Based on this, a mechanism where helix formation in ACTR occurs after accumulation of the initial encounter complex is unlikely. Therefore, the salt-sensitive kinetic phase λ^intermediate^ is probably not associated with any large overall changes in helicity. The model suggesting conformational selection in NCBD, and where all helices in ACTR are formed in the association step also agrees poorly with the experimental data. However, we do not completely rule out this model as this analysis was based on poorly defined rate constants from the fit of the fluorescence-monitored data to the conformational selection model (Figure 6c). The experimental data agrees best with an induced fit model, where all helical content is formed in the initial binding step(s) before accumulation of the intermediate. The subsequent conformational rearrangement step is thus likely not associated with large changes in helicity. Fitting of the stopped flow CD data directly to the induced fit model, using the rate constants in Figure 5 yielded values of the amplitude factors for [NCBD:ACTR] and [NCBD?ACTR]* of 5.26 ± 0.07 and 5.61 ± 0.05, respectively. The ratio between the amplitudes (5.26/ 5.61 ~ 0.94) suggests that more than 90 % helicity occurs in the association step. An induced fit model with helical intermediate is in excellent agreement with previous kinetic studies^8,39^ that favored induced fit models. Figure 7d summarizes our findings with an energy diagram for the best fitting model of induced fit with helical intermediates, where the conformational rearrangement in the final step is not associated with any significant change in helicity.

## Discussion

Coupled folding and binding reactions are mechanistically complex as they involve many interdependent structural changes that are challenging to monitor experimentally. A full mechanistic description of coupled folding and binding reactions thus necessarily involves integration of many different types of information. Here, we conducted the first rSRCD study of coupled folding and binding, and show that at least 90% of the helicity arises concomitant with association. This is similar to the nucleation-condensation mechanism in protein folding, where secondary and tertiary structure form simultaneously^40^. A main advantage of rSRCD is that the amplitude factors are structurally informative. The amplitude factors are sensitive to the structure in intermediates that accumulate during the reaction, and thus provide complementary information to fluorescence-monitored kinetics.

The rate-limiting step of the interaction between ACTR and NCBD has previously been probed by experiment and simulation^37,38,41^. The data suggested that the rate-limiting transition state occurs early in the reaction with partial native-like helix formation in for example helix 1 of ACTR and helix 3 of NCBD. Single-molecule studies with high time resolution suggests formation of an intermediate on the μs-time scale^21^, most likely corresponding to an electrostatically stabilized intermediate. This intermediate occurs on a much shorter timescale than can be detected by the rSRCD setup and is thus observed as part of the association phase. Our data suggests that once this energy barrier has been traversed, the remaining helices form rapidly and before the intermediate that accumulates at high ionic strength. Since we find that the helical content of the intermediate is native-like, it follows that the subsequent transition involves a rearrangement of folded helices. High ionic strength unmasks this step, so it is reasonable to assume that it involves electrostatic interactions. Electrostatics are important for initial association as shown by the salt dependence of the ACTR/NCBD interaction^42^. Previous studies have highlighted a buried ionic interactions as a determinant for packing specificity^13^, removal of which also has a strong negative effect on *k_on_*^8^. A compact helical complex with mproper ion pairing is thus a likely candidate for the kinetic intermediate observed at high ionic strength.

Coupled folding and binding reactions have been investigated thoroughly by molecular simulations^43^. The complex between NCBD and ACTR is one of the most popular model systems for such simulations. Some of these simulations have highlighted partially folded intermediates during the reactions. This prompts the question of whether the intermediates seen in simulations correspond to the intermediates giving rise to different kinetic phases? The measurement of helix formation in the different kinetic phases allows us to answer this question. The short answer is no. The intermediates reported in simulation studies all have an intermediate degree of helicity^23,24^. Since almost all helicity is gained in the first kinetic phase, this suggest any putative intermediates with partially formed helices would be occurring as very short-lived species upon barrier crossing. Simulations of coupled binding and folding reactions necessarily involve compromises in terms of coarse-graining, implicit water or various methods for enhanced sampling^43^. Such systems may not always reproduce the subtle energetic landscape of a coupled folding and binding reaction, and equivalence between intermediates observed in simulations and experiments should not be assumed automatically. The present study provides an experimental benchmark for future simulation studies regarding helix formation at different stages in the reaction.

The rSRCD facility presents a novel tool to interrogate structural changes accompanying biochemical reactions. The system studid here is challenging due to the rapid bimolecular binding kinetics with *k*_obs_ values around 100 s^-1^ and amplitudes with low signal-to-noise. We could only record reliable data within a narrow range of conditions, which suggests that we are at the edge of feasibility for the current setup. However, even with the present limitations the rSRCD setup is a promising new tool to follow binding reactions, especially for coupled folding and binding as such reactions often result in a change in CD signal. A key challenge for the continuing development of the rSRCD beamline is to reduce the instrumental artefact that obscures the first milliseconds. CD measurements are inherently more sensitive to the geometry of the cuvette than e.g. fluorescence, and therefore this problem may be hard to avoid completely, but will be addressed in further development of the facility.

## Acknowledgements

The rSRCD side branch was funded by the Carlsberg Foundation and the Danish Research Infrastructure Programme, and the AU-AMO beam line was additionally funded by the Lundbeck Foundation and the Independent Research Fund Denmark | Natural Sciences. Access to the beam line was granted by ISA, Centre for Storage Ring Facilities, Aarhus at Aarhus University. This work was supported by grants to M.K. from the “Young Investigator Program” of the Villum Foundation and the AIAS COFUND program (Agreement No. 609033) and by the Swedish Research Council to P.J. (grant 2016-04965). Lasse Staby and Mikhail Kuravsky are thanked for assistance with the calibration of the mixing time.

